# Observing intersubunit dynamics in single yeast ribosomes

**DOI:** 10.1101/2025.06.05.658109

**Authors:** Ananya Das, Amy K. Grove, Aleksandr V. Ivanov, Hironao Wakabayashi, Dmitri N. Ermolenko

## Abstract

Translation is accompanied by the rotation of the small and large ribosomal subunits relative to each other. Here, we use single-molecule Förster resonance energy transfer between fluorophores introduced into ribosomal proteins uS15 and eL30 to follow the intersubunit dynamics of *Saccharomyces cerevisiae* ribosomes. Similar to their bacterial counterparts, yeast ribosomes are observed to sample two predominant FRET states corresponding to the nonrotated (NR) and rotated (R) conformations. Our data yield further evidence that intersubunit rotation is coupled to tRNA transitions between the classical and hybrid binding states. In particular, the transition from the NR to R conformation is inhibited by the antibiotic cycloheximide, which is known to bind to the E site of the large subunit and prevent the movement of deacylated tRNA into the hybrid P/E state. The elongation cycle, which comprises tRNA binding, peptide transfer, and mRNA-tRNA translocation, is accompanied by switching from NR to R, and then back to the NR conformation. We find that fungal elongation factor 3 (eEF3) stabilizes the NR conformation of the ribosome. Our data are consistent with the model suggesting that eEF3 facilitates E-site tRNA release at the late step of mRNA-tRNA translocation, following the reverse intersubunit rotation induced by the universally conserved elongation factor 2 (eEF2). Our uS15-eL30 smFRET assay provides the basis for investigating eukaryotic mechanisms of translation regulation, including ribosome pausing, stalling, and frameshifting.

## INTRODUCTION

Protein synthesis is accompanied by large-scale, intricate conformational changes of the ribosome. These dramatic rearrangements include ∼10° rotation of the small and large ribosomal subunits relative to each other; ∼20° rotation of the head domain of the small subunit relative to its body, termed “head swivel”; a tilting motion of the head of the small subunit toward the large subunit in the direction orthogonal to head swivel; and inward/outward motion of the L1 stalk of the large subunit (1–3). In addition, the small subunit of the eukaryotic ribosome rotates around its elongated axis in a way that is orthogonal to the intersubunit rotation discovered in bacterial ribosomes (4,5). This motion was termed “subunit rolling” (4). Structural dynamics of the ribosome are essential for translation. For example, blocking intersubunit rotation by a disulfide crosslink between proteins on the small and large subunits was shown to abrogate ribosome translocation along mRNA (6).

In past decades, X-ray crystallography and cryo-EM have revealed dozens of snapshots of both bacterial and eukaryotic ribosomes that differ in degrees of intersubunit rotation, L1 stalk movement, swivel, and tilting of the head domain of the small subunit (1,7–9). However, elucidating the relative stability and prevalence of different structural intermediates requires real-time observation of ribosome dynamics in solution, free of averaging over asynchronous and heterogeneous populations of macromolecules. This gap in knowledge is filled with single-molecule Förster resonance energy transfer (smFRET) microscopy, which allows tracking of structural rearrangements by distance-dependent changes in energy transfer between single donor and acceptor fluorophores. Indeed, an extensive body of smFRET data has been generated to uncover the sequence and kinetics of rearrangements of the ribosome, tRNAs, and translation factors during different phases of bacterial translation (10–12).

In particular, smFRET data shows that during the translation elongation cycle, bacterial ribosomes adopt two predominant conformations: the nonrotated and fully rotated (13–15). Upon binding of the aminoacyl-tRNA to the A site followed by the peptidyl-transfer reaction, the ribosome transitions from the nonrotated into the rotated conformation. This rearrangement is coupled to the transition of peptidyl- and deacylated tRNAs from the classical A/A and P/P to the hybrid A/P and P/E states of tRNA binding (this nomenclature of binding sites denotes tRNA positions on the small and large ribosomal subunits respectively). tRNA and mRNA translocation on the small subunit is induced by the universally conserved ribosome-dependent GTPase (EF-G in bacteria or eEF2 in eukaryotes) and coincides with the reverse rotation of the ribosomal subunits back into the nonrotated conformation (10–12).

In contrast to studies of bacterial translation, only a few smFRET assays have been developed to examine structural rearrangements of the eukaryotic translation apparatus. In particular, tRNA binding and translocation in mammalian ribosomes was followed by smFRET between fluorophores introduced into the elbows of A- and P-site tRNAs (16–21). Fluorescent labeling of proteins uS19 and uL18 was employed to probe subunit joining during translation initiation in human and yeast ribosomes (22,23). However, these smFRET assays did not detect intersubunit rotation. To follow intersubunit rotation during non-canonical translation initiation on the cricket paralysis virus (CrPV) Internal Ribosomal Entry Site (IRES), fluorescently-labeled oligos were annealed to extensions engineered into helixes 44 and 101 of the 18S and 25S rRNAs in the small and large subunits of *Saccharomyces cerevisiae* ribosomes (24,25). Nevertheless, this assay is yet to be used to examine eukaryotic translation elongation. Therefore, the real-time conformational dynamics of eukaryotic ribosomes remain largely unexplored.

Here, we develop a single-molecule microscopy assay to examine intersubunit rotation coupled to the translation elongation cycle in yeast ribosomes. Our data reveal that intersubunit dynamics of yeast and bacterial ribosomes are similar. However, in contrast to bacterial pretranslocation ribosomes, which fluctuate between the nonrotated and rotated conformations with a frequency of 0.2-2 per second (13,15), yeast pretranslocation ribosomes rarely undergo apparent spontaneous intersubunit rotation.

In some fungi, including *S. cerevisiae*, eEF2-induced tRNA/mRNA translocation involves another elongation factor: an ATP-binding cassette (ABC) protein, elongation factor 3 (eEF3) (26–29). eEF3 is an essential gene in budding yeast (28). Ribo-seq analysis of translational changes induced by eEF3 depletion indicates that eEF3 is involved in the translocation step of the elongation cycle (27,28). Mutations in eEF3 also affect frameshifting efficiency (30). In ensemble kinetic experiments tracking incorporation of amino acids into the growing polypeptide chain, eEF3 accelerated the rate of tri- but not dipeptide formation, indicating that eEF3 is required to enable the second and, presumably, all following elongation cycles (28). Cryo-EM structures and biochemical experiments indicated that eEF3 binding leads to the opening of the L1 stalk and release of deacylated tRNA from the E site (26,28,29). Structural data also suggest that eEF3 may favor the non-rotated conformation of the ribosome (26,28), although eEF3 binding to the fully rotated ribosomes was also observed (5). Nevertheless, how eEF3 enables translation elongation and regulates structural rearrangements of the ribosome coupled to tRNA translocation has not been fully elucidated. In this work, we used our newly developed smFRET assay to further probe how eEF3 facilitates eEF2-induced translocation.

## METHODS

### Preparation of mRNAs and aminoacylated tRNAs

*E.coli* tRNA^Met^, tRNA^Val^ and tRNA^Tyr^, and *S. cerevisiae* tRNA^Phe^, were purchased from Chemblock. Bacterial and yeast tRNAs were aminoacylated using S100 cell extracts from *E. coli* and *S. cerevisiae*, respectively (31). Met-tRNA^Met^ was acetylated using acetic anhydride as previously described (31). The extent of aminoacylation was analyzed by acid gel electrophoresis (Supplementary Fig. S1) as previously described (32).

MFV (5ʹ G**AUG**UUUGUCUAUCAACAACAACAACAACAACAACAACAACAACAACAACAACAA<u>GGUUUUUCUUCUGAAGAUAAAG</u> 3ʹ) and MFY (5ʹ <u>GGUUUUUCUUCUGAAGAUAAAG</u>CAACAACAACAAGGCAAGGAGGUAAAA**AUG**UUCUACAA 3ʹ) were transcribed by T7 polymerase-catalyzed run-off transcription as previously described (33). The sequences complementary to the biotinylated AL2 DNA handle and the AUG start codon are <u>underlined</u> and **bold**, respectively.

### Purification of elongation factors

eEF1A, eEF1Bα, eEF2 and eEF3 were purified using previously published procedures with minor modifications (34–36).

Untagged eEF1A was purified from yeast strain BCY123 (37), which has the genotype *pep4::HIS3 prb::LEU2 bar1:HISG lys2::GAL1/10-GAL4 can1 ade2 ura3 leu2–3,112* and thus lacks several proteases. Cells grown to 0.9-1.1 OD_600_ were lysed by French press in 50 mL of lysis buffer containing 40 mM HEPES-KOH pH7.5, 50 mM KCl, 25% glycerol, 0.2 mM EDTA, with 0.2 mM PMSF and 1 tablet of Pierce protease inhibitor cocktail. Clarified lysate was loaded on 12.5 mL of DE52 resin. Column flowthrough was collected and then loaded on 3 mL of CM-sepharose. eEF1A was eluted by 10 mL of 40 mM HEPES-KOH pH 7.5, 300 mM KCl, 25% glycerol, 0.2 mM EDTA. The eluate was dialyzed against 40 mM HEPES-KOH pH 7.5, 50 mM KCl, 0.2 mM EDTA and loaded on UnoS6 cation exchange column (Bio-Rad) and eluted in 50-300 mM gradient of KCl concentration. Pure fractions were combined, concentrated via 10 kDa centricons (Millipore-Sigma) and stored in eEF storage buffer (50 mM Tris-HCl pH 7.5, 100 mM KCl, 10% glycerol and 1 mM DTT) at -80°C.

To purify eEF1Bα, BL21 *E.coli* cells were transformed with pET28b plasmid encoding eEF1Bα-TEV site-6His tag. eEF1Bα was expressed and purified via Ni-NTA resin (Thermo Scientific) using standard procedures (35). Cells were lysed in 50 mM Tris-HCl pH 7.5, 300 mM NaCl, 20 mM imidazole, 10% glycerol, 1 mM PMSF, and lysate was clarified via centrifugation before it was loaded on 2 mL Ni-NTA resin. Protein was eluted with buffer containing 250 mM imidazole. The 6His tag was cleaved off by TEV protease during overnight dialysis in 50 mM HEPES-KOH pH 7.5, 100 mM KCl, 10% glycerol, and then run over the nickel resin again in 10 mM imidazole. Purified protein was concentrated from the flowthrough and stored in eEF storage buffer (50 mM Tris-HCl pH 7.5, 100 mM KCl, 10% glycerol and 1 mM DTT) at -80°C.

TKY675 yeast strain expressing 6His-tagged eEF2 (34) was growth in 4L of YPD media to 1.0-4.0 OD_600_. Pelleted cells were lysed by French press in 50 mL of 50 mM Tris-HCl pH 7.5, 1M KCl, 5% glycerol, 20 mM imidazole, 0.025% tween-20, 0.5 mM PMSF and 1 tablet of Pierce protease inhibitor cocktail. After clarification, this lysate was loaded on 2 mL of Ni-NTA resin, which was subsequently washed with lysis buffer containing 40 mM imidazole. eEF2 was eluted by raising imidazole concentration to 250 mM. eEF2-containing fractions were dialyzed against 50 mM Tris-HCl pH 7.5, 200 mM KCl, 5% glycerol, 40 mM imidazole, loaded on Ni-NTA resin and eluted with 250 mM imidazole. eEF2 was concentrated using 10 kDa centricons (Millipore-Sigma, USA) and stored in 50 mM Tris-HCl pH 7.5, 200 mM KCl, 10% glycerol and 1 mM DTT at -80°C.

TKY1653 yeast strain expressing 6His-tagged eEF3 (36) was grown in YPD media. eEF3 was purified using Ni-NTA similar to eEF2. Subsequently, eEF3 was loaded onto UnoQ6 anion exchange column (Bio-Rad) and eluted in a 40-300 mM gradient of KCl concentration. Pure fractions were concentrated and stored in eEF storage buffer at -80°C.

### ybbR tagging of protein uS15 and eL30

The sequence encoding a C-terminal ybbR peptide (DSLEFIASKLA) with triple glycine linkers at both ends of the ybbR peptide (GGG-ybbR-GGG) was introduced into *RPS13* (uS15) and *RPL30* (eL30) according to the method described (38). BCY123 cells were transformed with a DNA cassette containing the uS15 or eL30 ORF, ybbR encoding sequence and HygrB DNA flanked by ∼1000 nt-long sequences homologous to the sequences flanking the respective ORFs in the yeast genome. The HygrB DNA encodes bacterial hygromycin B phosphotransferase, which produces resistance to the antibiotic hygromycin B. These DNA constructs were created by Gibson assembly (39) and verified by sequencing. To prepare competent yeast cells for transformation, yeast cells were growth in YPD liquid media to OD_600_ between 1.3 and 2.5. For each transformation, 10 mL of cell culture was pelleted by centrifugation at 3000 rpm for 5 minutes. Pellets were washed twice with 1 mL of 100 mM lithium acetate. Next, cells were resuspended in 110 μl of 100 mM lithium acetate containing salmon sperm DNA at concentration of 1 mg/mL, and then incubated with 300 ng of linearized DNA at 30°C for 15 minutes. Next, 600 μl of a lithium acetate-PEG solution (48% PEG 3350, 0.1 M lithium acetate) was added. After incubation at 30°C for 30 minutes, cells were mixed with 68 μl of DMSO and incubated at 42°C in for 15 minutes. After transformation, cells were pelleted at 1500 rpm for 7 minutes, resuspended in 1 mL of YPD media and incubated at 30°C for 1 hour while shaking. After this recovery period, 250 μl of cells were grown on a YPD-HygrB plate containing hygromycin B (50 μg/mL) at 30°C for 2 days. Individual colonies were streaked onto another YPD-HygrB plate and grown at 30°C for 2 days. Mutations were verified by sequencing of the respective ORFs in the genomic DNA.

### Purification of 40S and 60S ribosomal subunits

Yeast ribosomal subunits were purified according to published procedures (40) with minor modifications. Yeast cells were cultured in 2 L of YPD media with 50 μg/mL hygromycin B up to OD600 0.9-1.1. Cells were pelleted at 12,000 g for 12 minutes at 4°C. The pellets were washed by 50 mL of ice-cold lysis buffer containing 20 mM Tris-HCl pH 7.5, 50 mM KCl, 10 mM MgCl_2_, 0.1 mM EDTA. Cells were lysed by French press in 35 mL of lysis buffer supplied with one tablet of Pierce protease inhibitor cocktail and heparin (1 mg/mL). The lysate was clarified by centrifugation at 14,000 rpm for 30 min and layered on 5 mL of lysis buffer containing 1.1 M sucrose. Ribosomes were pelleted by centrifugation with a Ti-70 rotor (Beckman Coulter) at 46,000 rpm for 21 hours and resuspended in 20 mM HEPES-KOH pH 7.5, 50 mM KCl, 4 mM MgCl_2_, 0.2 mM EDTA and 8.55% sucrose. KCl concentration was adjusted in a drop-wise manner to 500 mM and then ribosomes were pelleted again by centrifugation in Ti-70 rotor. Ribosomes were resuspended in 40S-60S buffer (20 mM HEPES-KOH pH 7.5, 500 mM KCl, 4 mM MgCl_2_, 0.2 mM EDTA) containing 10 mM puromycin and incubated for 10 minutes at 30°C. 100-300 units of OD_260_ were layered on 10-30% sucrose gradient prepared in 40S-60S buffer and centrifuged with a SW-28 rotor (Beckman Coulter) at 20,000 rpm for 18 hours. 40S and 60S fractions were concentrated by 100 kDa centricons (Millipore-Sigma, USA) and stored in 50 mM HEPES-KOH pH 7.5, 50 mM KCl, 2.5 mM MgCl2, 0.2 mM EDTA, 8.55% sucrose at -80°C.

### Ribosome labeling

40S with ybbR-tagged uS15 and 60S with ybbR-tagged eL30 were conjugated by Sfp phosphopantetheinyl transferase (41) to Cy3-CoA and Cy5-CoA, respectively. To express Sfp, *E.coli* BL21 cells were transformed with pET29b vector encoding C-terminal 6His tag-Sfp. The Sfp pET29b plasmid was a gift from Michael Burkart (Addgene plasmid # 75015 ; http://n2t.net/addgene:75015; RRID:Addgene_75015) (42). Sfp was purified on Ni-NTA resin using standard procedures. Transformed BL21 cells were grown in liquid culture at 37°C until OD_600_ = 0.6, and then induced with 200 mM IPTG and shaken at 18°C overnight. Cells were harvested via centrifugation and lysed via French press. Cell debris was precipitated via centrifugation, and then the supernatant was passed through Ni-NTA resin equilibrated in lysis buffer (50 mM NaH_2_PO_4_, 300 mM NaCl, 5 mM imidazole; pH 8). The resin was washed with wash buffer (50 mM NaH_2_PO_4_, 300 mM NaCl, 20 mM imidazole; pH 8), and then Sfp was eluted with 50 mM NaH_2_PO_4_, 300 mM NaCl, 200 mM imidazole, pH 8. After SDS-PAGE analysis, the protein was concentrated and stored at -80°C in 50 mM Tris-HCl pH 7.5, 10mM MgCl_2_ until use.

100 μM coenzyme A tri-lithium salt (Sigma-Aldrich) was incubated with either 200 μM Cy3-malemide or Cy5-maleimide (both from Click Chemistry Tools) in 100 mM sodium phosphate, pH 7.0 for one hour at room temperature. Reaction was stopped by 2 mM β-mercaptoethanol. Next, 20 μM fluorophore-CoA conjugate was incubated with 10 μM ybbR-tagged ribosomal subunits and 20 μM Sfp phosphopantetheinyl transferase in 20 mM HEPES, pH 7.6, 0.5% Tween 20, 200 mM KCl, 10 mM MgCl_2_, 0.25 M sucrose for 1 hour at 30°C. Labeled subunits were layered onto a 10-30% sucrose gradient containing 20 mM HEPES-KOH pH 7.5, 500 mM KCl, 4 mM MgCl_2_, 0.2 mM EDTA, and centrifuged in a SW-41 rotor at 20,000 rpm for 18 hours. Ribosomal fractions were concentrated by 100 kDa centricons (Millipore-Sigma, USA) and stored in 50 mM HEPES-KOH pH 7.5, 50 mM KCl, 2.5 mM MgCl_2_, 0.2 mM EDTA, 8.55% sucrose at -80°C. To produce uS15-Cy3/eL30-Cy5 80S ribosomes, labeled 40S and 60S subunits were associated in 50 mM HEPES-KOH pH 7.5, 50 mM KCl, 4 mM MgCl_2_, 3.5 mM spermidine, 0.2 mM EDTA, 8.55% sucrose at 30°C for 20 minutes and subsequently purified by centrifugation in a 10-30% sucrose gradient.

### Assembly of ribosomal complexes

All ribosomal complexes were assembled in ribosome buffer containing 50 mM HEPES-KOH pH 7.5, 50 mM KCl, 2 mM spermidine, 2.5 mM MgCl_2_, 6 mM β-mercaptoethanol and 0.025% non-ionic detergent Nikkol (octaethylene glycol monododecyl ether, Sigma-Aldrich).

The 3ʹ biotinylated AL2 DNA oligo (5ʹ CTTTATCTTCAGAAGAAAAACC-biotin 3ʹ, IDT) was annealed to MFY mRNA via 3-minute incubation at 66°C followed by placing mRNA/DNA sample on ice for 5 minutes. Likewise, the 5ʹ biotinylated AL2 oligo (IDT) was annealed to leaderless MFV mRNA. To fill the ribosome P site with peptidyl-tRNA analogue N-Ac-Met-tRNA^Met^, 0.3 μM uS15-Cy3/eL30-Cy5 80S ribosomes were incubated with 0.6 μM mRNA-AL2 and 0.6 μM N-Ac-Met-tRNA^Met^ for 20 minutes at 30°C. To assemble the aa-tRNA-eEF1A-GTP ternary complex, 7.2 μM eEF1A, 10.9 μM eEF1Bα and 3.3 mM GTP/MgCl_2_ were incubated for 20 minutes at 30°C. Next, 3.7 μM aa-tRNA (Phe-tRNA^Phe^, Val-tRNA^Val^ or Tyr-tRNA^Tyr^) was added, followed by a 10-minute incubation at 30°C. Then 0.5 μM aa-tRNA-eEF1A-GTP ternary complex was incubated with 0.26 μM 80S containing P-site N-Ac-Met-tRNA^Met^ for 15 minutes at 30°C. To induce translocation, 0.18 μM pretranslocation 80S ribosomes were incubated with 1 μM eEF2 and 0.5 mM GTP/MgCl_2_ in the absence or presence of 1 μM eEF3 and 0.5 mM ATP/MgCl_2_ for 10 minutes at 30°C.

### smFRET measurements

Cy3 was excited by a 532 nm laser; Cy3 and Cy5 fluorescence was measured by a home-built prism-type inverted TIRF (Total Internal Reflection Fluorescence) microscope equipped with a water-immersion objective (Olympus, UplanApo, 60x, NA 1.2), and a Hamamatsu Fusion Back Illuminated sCMOS camera and Gemini Dual Image Splitter, which uses a 630-nm dichroic mirror to parse fluorescence emission into Cy3 and Cy5 detection channels. Quartz slides were coated with dichlorodimethylsilane (DDS, Fisher), bound with biotin-BSA (Fisher) and passivated by Tween-20 (Fisher) as previously described (43). To enable tethering to biotin-BSA, 40 μL 0.2 mg/mL neutravidin (Fisher) was added to the slide in H50 buffer (10 mM HEPES, pH 7.5 and 50 mM KCl). Non-specific sample binding to the slide was checked in the absence of neutravidin. Ribosomal complexes were diluted to 4 nM with the ribosome buffer containing 0.025% Nikkol and added to the slide. To image immobilized ribosomes, unbound ribosomes were washed off and the slide chamber was filled with 50 μl of imaging buffer containing 50 mM HEPES-KOH pH 7.5, 50 mM KCl, 2 mM spermidine, 2.5 mM MgCl_2_, 6 mM beta-mercaptoethanol, 0.8 mg/mL glucose oxidase, 0.625% glucose, 0.4 μg/mL catalase and 1.5 mM 6-hydroxy-2,5,7,8-tetramethylchromane-2-carboxylic acid (Trolox, Sigma-Aldrich). In kinetic experiments (Fig. 7), 50 μl of imaging buffer with 1 μM eEF2, 0.4 mM GTP-MgCl_2_, 1 μM eEF3, and 0.4 mM ATP-MgCl_2_ (if translocation was performed in the presence of both eEF2 and eEF3) was injected into the slide with immobilized pretranslocation ribosomes using a J-Kem syringe pump.

3-minute-long movies were acquired using the Micromanager open-source software (https://micro-manager.org/) with 100 ms integration time. All smFRET measurements were performed at room temperature (22 °C). Fluorescence intensities of Cy3 donor (*I_D_*) and Cy5 acceptor (*I_A_*), from single ribosomes were extracted via the vbscope software (https://github.com/GonzalezBiophysicsLab/vbscope-paper). Apparent FRET efficiency (E *_FRET_*, hence referred as FRET) was calculated as follows:

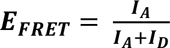

smFRET data were analyzed using the tMAVEN software (44) (https://gonzalezbiophysicslab.github.io/tmaven/). FRET distribution histograms compiled from hundreds to thousands of smFRET traces were plotted with 0.02 binning size and fit to two Gaussians corresponding to 0.52 (mean) ±0.01 (s.d.) and 0.80 (mean) ±0.02 (s.d.) FRET in OriginLab (Supplementary Table S1). To analyze FRET transitions, smFRET traces were idealized by 2-state Hidden Markov Model using an ebFRET algorithm (45) in tMAVEN (44). The rate of reverse intersubunit rotation coupled to translocation was determined by fitting the distribution of τ_trl_ to a single exponential decay function.

## RESULTS AND DISCUSSION

### Fluorescent labeling of yeast ribosomal subunits

To follow intersubunit rearrangements in yeast ribosomes, we labeled ribosome proteins uS15 and eL30 with a donor (Cy3) and an acceptor (Cy5) fluorophores, respectively, via a ybbR tagging strategy (46). This approach was previously successfully used for labeling of ribosomes and translation factors (22,23,47). Both proteins are largely solvent-exposed and located peripherally relative to the intersubunit interface (Fig. 1). uS15 and eL30 are also distant from the binding sites of translation factors and tRNAs (48). Hence, modifying proteins uS15 and eL30 is unlikely to perturb ribosome function. Unlike many ribosomal proteins in yeast, which are encoded by two paralogues, uS15 and eL30 are encoded by single genes *RPS13* and *RPL30,* which facilitates genetic manipulations for tagging.

**Figure 1.**
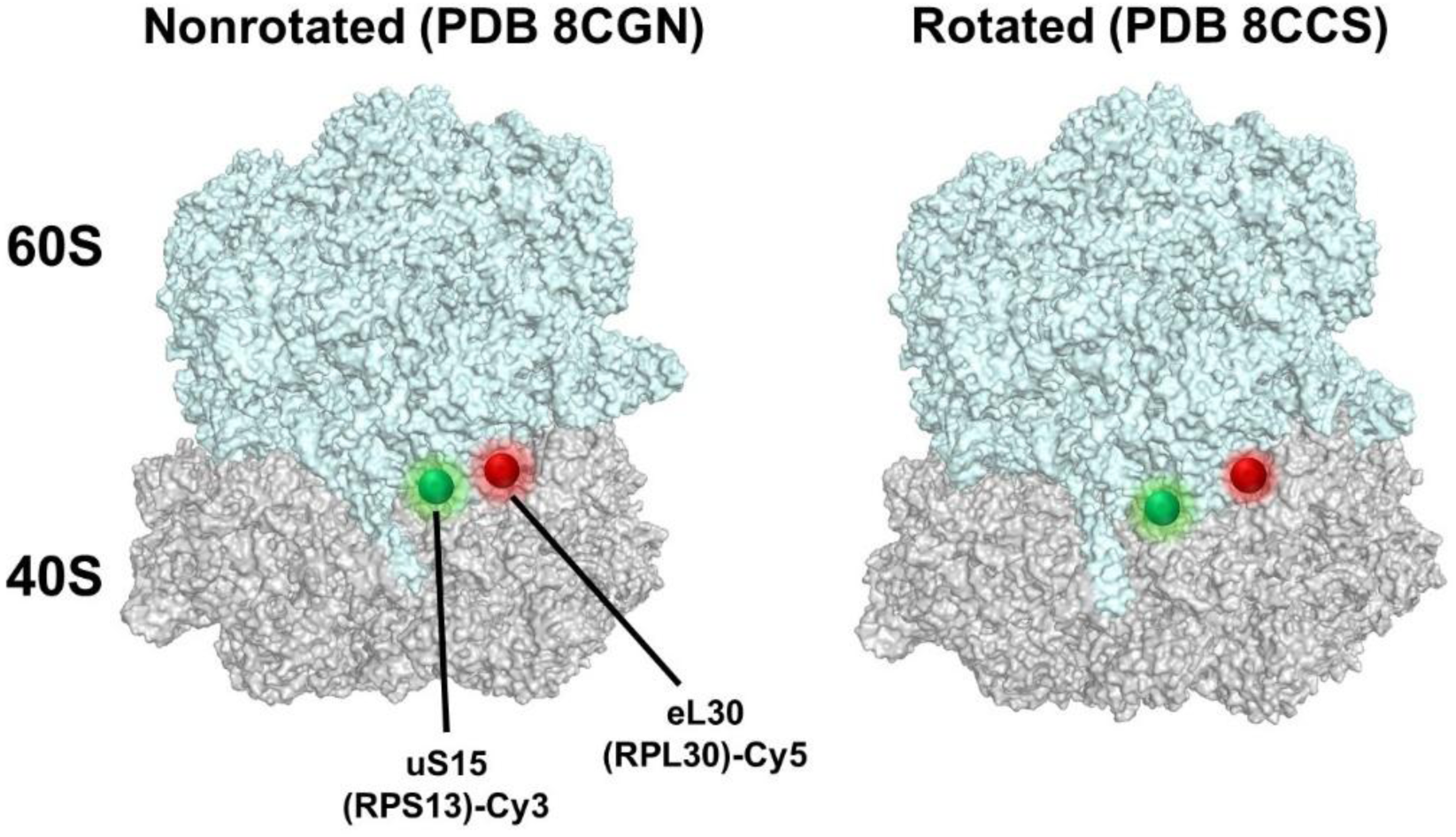
Introduction of donor and acceptor fluorophores into yeast ribosomes. Donor (Cy3, green sphere) and acceptor (Cy5, red sphere) fluorophores were conjugated to the C termini of proteins uS15 and eL30 on the small (grey) and large (cyan) ribosomal subunits of the yeast 80S ribosome. Cryo-EM structures of the nonrotated, posttranslocation (PDB 8CGN) and fully-rotated, pretranslocation (PDB 8CCS) yeast ribosomes (49) aligned by 25S rRNAs show that the distance between the C-termini of uS15 and eL30 changes from 28 Å (nonrotated) to 35 Å (rotated).

An 11 amino acid-long ybbR tag, together with a hygromycin B resistance cassette (*ybbR:HygrB*), was introduced at the C-terminus of either uS15 or eL30 in the genetic background of BCY123 yeast strain (37) via homologous recombination. The first serine residue of the ybbR tag can be enzymatically linked to the fluorophore-derivatized phosphopantetheinyl group of co-enzyme A via Sfp transferase (46). The BCY123 strain lacks several proteases and, thus, is particularly suitable for ribosome purification (37). The distances between C-terminal carbonyl oxygen atoms in uS15 (Asn 151) and eL30 (Ala 105) measured in cryo-EM structures of the fully-rotated, pretranslocation (PDB 8CCS) and nonrotated, posttranslocation (PDB 8CGN) yeast ribosomes were 35 and 28 Å, respectively (49). The ybbR tag and flexible glycine linker is expected to extend these distances and bring them closer to the R_0_ of a Cy3-Cy5 pair (56 Å) (50) where FRET is most sensitive to changes in inter-fluorophore spacing. Hence, donor-acceptor FRET pair introduced at the C termini of uS15 and eL30 should be sensitive to intersubunit rotation.

The *RPL30-ybbR:HygrB* strain grew as well as the wild-type (BCY123) yeast cells while growth of the *RPS13-ybbR:HygB* strain was only slightly attenuated (Supplementary Fig. S2A). Therefore, C-terminal ybbR tagging of either eL30 or uS15 did not substantially affected ribosome function or biogenesis. We next purified the small and large ribosomal subunits from *RPS13-ybbR:HygrB* (uS15) and *RPL30-ybbR:HygrB* (eL30) strains and labeled them with Cy3 and Cy5 fluorophores, respectively. SDS-PAGE analysis indicated high specificity of labeling as proteins uS15 and eL30 were conjugated with Cy3 and Cy5 fluorophores, respectively (Supplementary Fig. S2B). Efficiency of labeling for both proteins varied among different preparations between 50 and 100%. After labeling, we associated 40S and 60S ribosomal subunits and purified 80S ribosomes via centrifugation in a sucrose gradient. We then used either ribosomal subunits or 80S ribosomes for elongation complex assembly.

### Non-enzymatic assembly of translation elongation complexes

Although translation initiation requires cooperative action of numerous initiation factors, complexes bound with tRNAs and capable of transitioning through all steps of the elongation cycle can be assembled in the absence of initiation factors with both bacterial and eukaryotic ribosomes. Because of its high affinity, the P site of the ribosome can be non-enzymatically loaded with tRNA that positions a cognate mRNA codon in the P site and defines the reading frame. This strategy was successfully used for assembly of eukaryotic ribosomal complexes in the absence of initiation factors for cryo-EM and smFRET experiments (16,51,52).

To non-enzymatically fill the P site, we incubated uS15-Cy3/eL30-Cy5 80S ribosomes with peptidyl-tRNA analogue N-Ac(acetyl)-Met-tRNA^Met^ in the presence of leaderless MFV mRNA, which contains a 5ʹ guanine base upstream of an AUG codon. Ribosomal complexes were then tethered to the microscope slide via a biotinylated DNA oligo annealed at the 3ʹ end of the mRNA (Fig. 2A-B). To optimize complex assembly, we tested several buffer conditions previously described in other smFRET and biochemical studies of eukaryotic ribosomes (17,23,24,40). The largest number of mRNA-bound uS15-Cy3/eL30-Cy5 80S ribosomes on the slide was reproducibly observed when ribosomal complex assembly was carried out in the presence of 2.5 mM MgCl_2_, 50 mM KCl and 2 mM spermidine (i.e., in a slightly modified buffer system used by Pestova and co-workers (40)). Increasing Mg^2+^ concentration to 5 mM did not substantially affect the number of observed smFRET traces, while raising KCl concentration to 100 mM reduced the number of smFRET traces ∼4-fold, indicating lessened stability of ribosome complexes. In 2.5 mM MgCl_2_, 50 mM KCl and 2 mM spermidine, we observed a negligible number of mRNA-bound 80S ribosomes on the slide if they were originally incubated with mRNA in the absence of tRNA or in the presence of deacylated tRNA^Met^. Therefore, the peptidyl moiety of the P-site tRNA is critical for stabilizing 80S ribosomal complexes.

**Figure 2.**
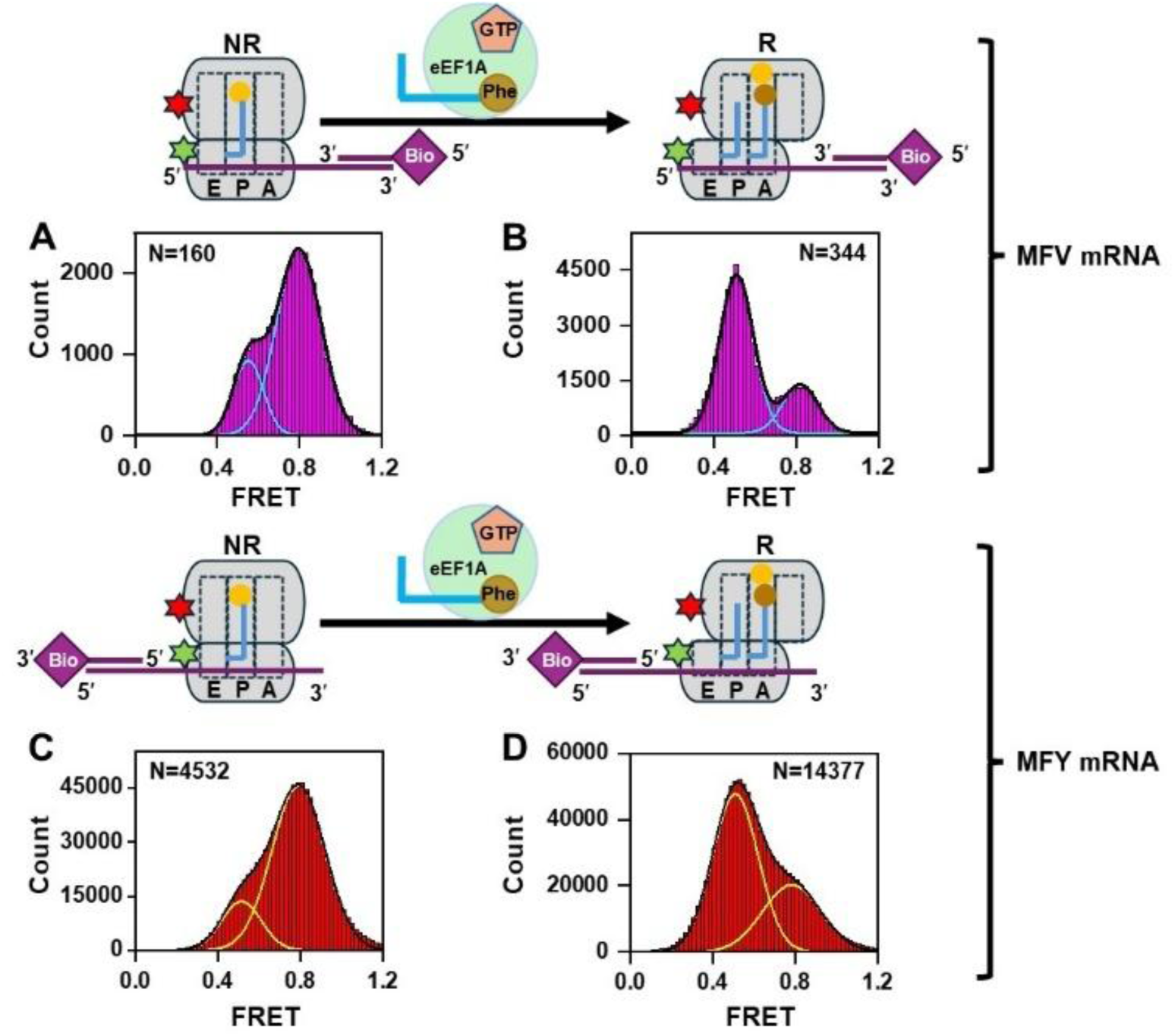
Binding of aa-tRNA to the A site, followed by peptidyl transfer, induces switching from the NR (0.8 FRET) to R (0.5 FRET) conformation of the ribosome. Histograms (A-D) show FRET distribution in uS15-Cy3/eL30-Cy5 ribosomes programmed with either leaderless, 3′ tethered MFV mRNA (A-B) or 5′ tethered MFY mRNA (C-D). Ribosomal complexes were immobilized on the microscope slide via a biotinylated DNA handle annealed to mRNA, as indicated by the schematics. Ribosomes bound with P-site N-Ac-Met-tRNA^Met^ (A, C) were incubated with 1 μM eEF1A•GTP•Phe-tRNA^Phe^ (B, D). Cyan and yellow lines indicate individual Gaussian fits. The black line shows the sum of Gaussian fits. N indicates the number of FRET traces compiled into each histogram.

The majority (80%) of uS15-Cy3/eL30-Cy5 80S ribosomes bound with P-site N-Ac-Met-tRNA^Met^ showed 0.8 FRET, while in the remaining 20% of ribosomes, 0.5 FRET was observed (Fig. 2A). Based on cryo-EM reconstructions of yeast ribosomes (49,52), intersubunit rotation is expected to increase the distance between the C-termini of uS15 and eL30. Therefore, 0.8 and 0.5 FRET likely correspond to the nonrotated and rotated conformations, respectively. Our smFRET data indicate that, consistent with previous structural and smFRET investigations of bacterial ribosomes (10,53), peptidyl-tRNA bound in the P site stabilizes the nonrotated conformation of eukaryotic ribosomes (0.8 FRET). Although complexes with deacylated P-site tRNA were observed in our experiments to be less stable than ribosomes with P-site peptidyl-tRNA, we cannot exclude the presence of ribosomes with deacylated tRNA bound in the hybrid P/E state that give rise to a low-FRET state population. It is also possible that because of the short N-Ac-Met moiety, N-Ac-Met-tRNA^Met^ is not completely fixed in the classical P/P state, and thus a minor fraction of ribosomes can adopt the rotated state. Finally, tRNA movements and intersubunit rotation may be correlated but not strictly coupled. Indeed, molecular dynamics simulation analysis indicated that while NR-to-R transition disfavors tRNA binding in the classical P/P state, the R conformation of the ribosome is compatible with both hybrid P/E and classical P/P states of tRNA binding (54).

### Yeast ribosomes switch between nonrotated and rotated conformations

To explore intersubunit dynamics during the elongation cycle, uS15-Cy3/eL30-Cy5 80S ribosomes bound with P-site N-Ac-Met-tRNA^Met^ were incubated with eEF1A•GTP•Phe-tRNA^Phe^ ternary complex and then imaged. eEF1A-mediated binding of aminoacyl-tRNA to the A site, followed by peptidyl transfer from P- to A-site tRNA, increased the fraction of ribosomes showing 0.5 FRET from 20 to ∼80% (Fig. 2B). This is consistent with transitioning from the nonrotated, classical into the rotated, hybrid state conformation of the ribosome bound with A/P-site N-Ac-Met-Phe-tRNA^Phe^ and P/E-site tRNA^Met^ (Fig. 2).

We next tested whether non-enzymatic ribosome complex assembly is restricted to the 5ʹ end of mRNA or also possible at internal mRNA codons. To this end, we incubated uS15-Cy3/eL30-Cy5 80S ribosomes with the peptidyl-tRNA analogue N-Ac-Met-tRNA^Met^ in the presence of MFY mRNA, which contains a unique AUG codon 49 nucleotides downstream of the 5ʹ end. Ribosomal complexes were then tethered to the microscope slide via a biotinylated DNA oligo, which was annealed at the 5ʹ end of the mRNA before the incubation with ribosomes (Fig. 2C-D). Similar to the P-site N-Ac-Met-tRNA^Met^-bound ribosomes programmed with leaderless MFV mRNA, ∼80% of ribosomes bound to MFY mRNA showed 0.8 FRET (Fig. 2C). Interestingly, when the same concentration of ribosomes was added to the slide, we consistently observed twice as many single-molecule traces per movie in complexes assembled with MFY mRNA than in ribosomes programmed with leaderless MFV mRNA, which indicates higher stability of the former. The 5’ leader sequence upstream of the start codon likely stabilizes the mRNA-ribosome complex by creating more extensive interactions with the mRNA channel on the small subunit.

Because of higher complex stability, we performed most of the following experiments with ribosomes bound to 5ʹ-tethered MFY mRNA. Evidently, non-enzymatic assembly of yeast ribosomal complexes can effectively occur on codons distant from the 5ʹ end and is not hindered by the biotinylated DNA oligo annealed at the 5ʹ end. Hence, 80S ribosomes can either directly bind to internal codons or bind to the 3ʹ end and slide over the mRNA in a 3ʹ-to-5ʹ direction.

Next, uS15-Cy3/eL30-Cy5 80S ribosomes programmed with MFY mRNA and bound with P-site N-Ac-Met-tRNA^Met^ were incubated with the eEF1A•GTP•Phe-tRNA^Phe^ ternary complex and then imaged. Phe-tRNA^Phe^ binding to the A site followed by peptidyl transfer reaction triggered switching from 0.8 to 0.5 FRET, similar to experiments performed with ribosomes bound to leaderless MFV mRNA (Fig. 2D).

We next examined how antibiotic didemnin B, which binds to eEF1A and traps eEF1A -aa-tRNA at the intermediate step of tRNA accommodation (55), affects the switching from NR to R conformation. When eEF1A•GTP•Phe-tRNA^Phe^ was incubated with didemnin B prior to incubation with ribosomes bound to P-site tRNA (Fig. 3), the ribosomes predominantly remained in the 0.8 FRET state corresponding to the NR conformation. Similar results were obtained when GTP was omitted during ternary complex assembly and the ribosome was incubated with just eEF1A and Phe-tRNA^Phe^ (Supplementary Fig. S3). Hence, eEF1A-mediated Phe-tRNA^Phe^ binding and peptidyl transfer are required for the transition from NR to R conformation of the ribosome.

**Figure 3.**
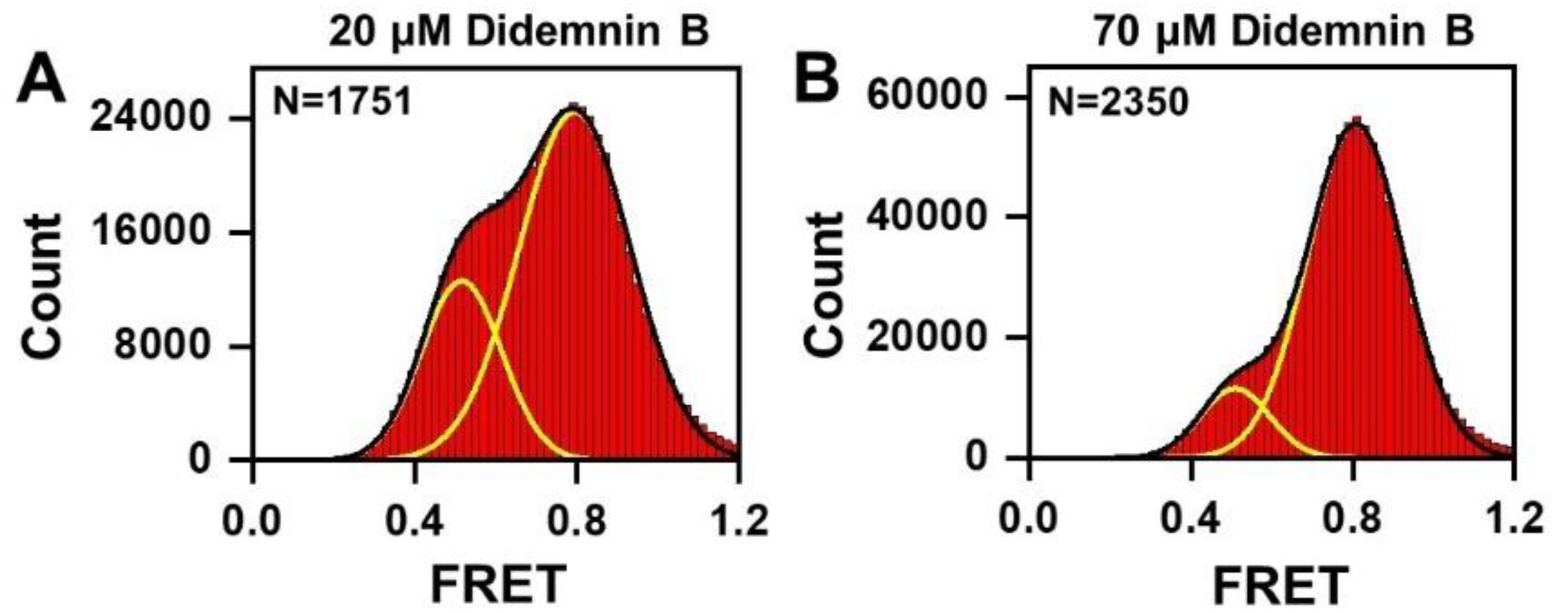
Didemnin B prevents eEF1A-mediated tRNA binding and switching of the ribosome from the NR to R conformation. eEF1A, eEF1B, and GTP were pre-incubated with 20 (A) and 70 μM (B) concentrations of the eEF1A inhibitor didemnin B (55) before incubation with Phe-tRNA^Phe^ and subsequent incubation with 80S ribosomes bound to MFY mRNA and P-site N-Ac-Met-tRNA^Met^. Yellow and black lines indicate individual Gaussian fits and the sum of Gaussian fits, respectively. N indicates the number of FRET traces compiled into each histogram.

We observed similar switching from 0.8 to 0.5 FRET upon incubation of eEF1A•GTP•Phe-tRNA^Phe^ with ribosomes bound with P-site N-Ac-Met-tRNA^Met^ when ribosomal complexes were assembled on MFY mRNA from the 40S and 60S subunits instead of 80S ribosomes (Supplementary Fig. S4A-B). However, significantly fewer smFRET traces per movie were detected for this assembly method, indicating the higher efficiency of non-enzymatic mRNA/tRNA loading of 80S ribosomes.

Our FRET measurements in uS15-Cy3/eL30-Cy5 80S ribosomes indicate that, akin to their bacterial counterparts, eukaryotic ribosomes sample two predominant conformations. Upon A-site tRNA binding and peptidyl transfer, yeast ribosomes transition from the nonrotated to rotated conformation.

### Intersubunit rotation in eukaryotic ribosomes is coupled to tRNA movement between the classical and hybrid binding states

Structural and smFRET studies revealed that in the absence of a catalyst of translocation (elongation factor G (EF-G)), bacterial pretranslocation ribosomes containing A-site peptidyl and P-site deacylated tRNAs spontaneously fluctuate between the nonrotated and rotated conformations (13,15). This spontaneous intersubunit rotation is thought to be coupled to inward/outward movement of the L1 stalk and tRNA fluctuations between the classical and hybrid binding states (10). Previously, yeast ribosomes bound to CrPV IRES and eEF2 trapped on the ribosome by the antibiotic sordarin were also found to undergo spontaneous cyclic NR↔R transitions (24). By contrast, pretranslocation uS15-Cy3/eL30-Cy5 80S ribosomes imaged in the absence of eEF2 remained static in either a 0.5 or 0.8 FRET state (Fig. 4A-B). Only ∼2% of traces in pretranslocation uS15-Cy3/eL30-Cy5 80S ribosomes bound with N-Ac-Met-Phe-tRNA^Phe^ and deacylated tRNA^Met^ exhibited spontaneous fluctuations between 0.5 and 0.8 FRET (Fig. 4C). We also did not observe the presence of traces showing intermediate FRET values, which could be indicative of averaging over frequent fluctuations between 0.5 and 0.8 FRET occurring faster than the 100 ms time resolution of our measurements. However, we cannot exclude the possibility that spontaneous fluctuations between the nonrotated and rotated conformations occur at rates much faster than the 100 ms time resolution of our smFRET experiments. Several experimental parameters, such as buffer conditions and N-acetyl moiety at the A-site N-Ac-Met-Phe-tRNA^Phe^, may contribute to reducing the rate of spontaneous intersubunit rotation in our experiments (15,56–58). Nevertheless, yeast pretranslocation ribosomes appear to be appreciably less dynamic than their bacterial counterparts.

**Figure 4.**
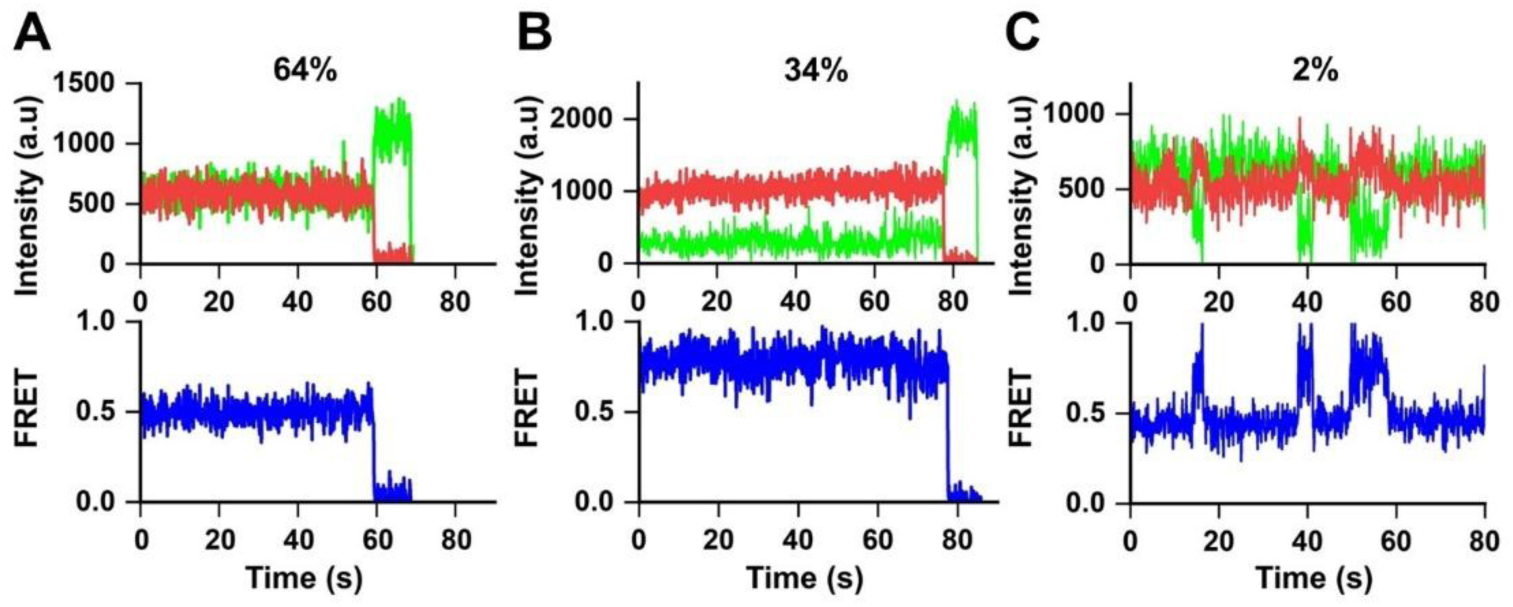
uS15-Cy3/eL30-Cy5 pretranslocation ribosomes sample 0.5 (R) and 0.8 (NR) FRET states. smFRET traces in pretranslocation ribosomes, which were bound with MFY mRNA, A-site N-Ac-Met-Phe-tRNA^Phe^ and P-site deacylated tRNA^Met^, show fluorescent intensities of Cy3 (green) and Cy5 (red) as well as FRET efficiency (blue). Most traces appear static either in the 0.5 FRET (64% traces, A) or 0.8 (34% traces, B) FRET states. ∼2% of traces show spontaneous fluctuations between 0.5 (R) and 0.8 (NR) FRET states (C).

To explore the connection between intersubunit rotation and tRNA movements into the A/P and P/E hybrid states, pretranslocation uS15-Cy3/eL30-Cy5 80S ribosomes bound with N-Ac-Met-Phe-tRNA^Phe^ and deacylated tRNA^Met^ were incubated with various concentrations of the eukaryotic antibiotic cycloheximide. Cycloheximide binds to the E site of the large subunit and thus hinders the movement of deacylated tRNA into the hybrid P/E state (59). smFRET experiments performed in pretranslocation yeast and mammalian ribosomes have previously demonstrated that cycloheximide stabilizes fluorescently labeled tRNAs in the classical A/A and P/P binding sites (17,21). Our measurements revealed that cycloheximide converts most of our pretranslocation uS15-Cy3/eL30-Cy5 80S ribosomes from a rotated to nonrotated conformation in a concentration-dependent manner (Fig. 5). Hence, intersubunit rotation in yeast ribosomes is coupled to tRNA movements between the classical and hybrid states of tRNA binding.

**Figure 5.**
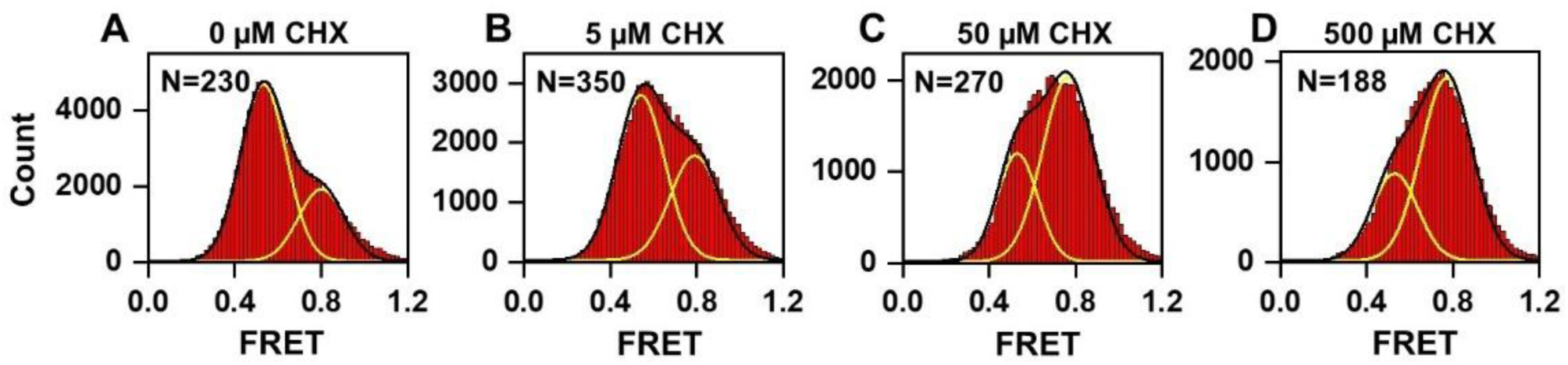
The antibiotic cycloheximide (CHX) stabilizes the NR conformation (0.8 FRET) in pre-translation ribosomes. Histograms (A-D) show FRET distribution in pretranslocation uS15-Cy3/eL30-Cy5 ribosomes incubated with different concentrations of CHX as indicated. Yellow and black lines indicate individual Gaussian fits and the sum of Gaussian fits, respectively. N indicates the number of FRET traces compiled into each histogram.

### The eEF3-mediated step in mRNA-tRNA translocation follows reverse intersubunit rotation

Following tRNA binding and peptide transfer, the translation elongation cycle is completed by mRNA-tRNA translocation induced by EF-G•GTP in bacteria or eEF2•GTP in eukaryotes. In bacteria, translocation of peptidyl- and deacylated tRNAs with their associated codons from the A and P to P and E sites of the small subunits was shown to be coupled to the transition from the rotated to nonrotated conformation of the ribosome (13,14,60). We tested whether eEF2-induced translocation in eukaryotic ribosomes is also linked to reverse intersubunit rotation and whether fungal elongation factor eEF3 affects the extent of translocation over two translocation cycles (Supplementary Fig. S5).

When we incubated pretranslocation uS15-Cy3/eL30-Cy5 80S ribosomes, which contained A-site N-Ac-Met-Phe-tRNA^Phe^ and P-site deacylated tRNA^Met^, with eEF2•GTP for 10 minutes at 30°C before imaging, the fraction of ribosomes in the nonrotated (0.8 FRET) conformation increased from ∼35 to ∼50%, consistent with coupling of mRNA-tRNA translocation to reverse intersubunit rotation (Fig. 6A, C). Incubating pretranslocation uS15-Cy3/eL30-Cy5 80S with both eEF2•GTP and eEF3•ATP reproducibly led to a larger shift in NR-R ratio as the fraction of ribosomes in the nonrotated (0.8 FRET) conformation increased to ∼60% (Fig. 6B, C). eEF3 is thought to be required to complete translocation by removing the deacylated E-site tRNA from the ribosome that affects the rate of the next elongation cycle (28). However, the Phe-tRNA^Phe^ used to load the A site during ribosome complex assembly contains deacylated tRNA (Supplementary Fig. S1), which may bind to the E site of pretranslocation ribosomes. eEF3-facilitated dissociation of this E-site tRNA likely increases the fraction of pretranslocation ribosomes undergoing translocation during the first elongation cycle in our *in vitro* experiments. These results are consistent with our eEF3 preparations being active in promoting eEF2-induced mRNA-tRNA translocation.

**Figure 6.**
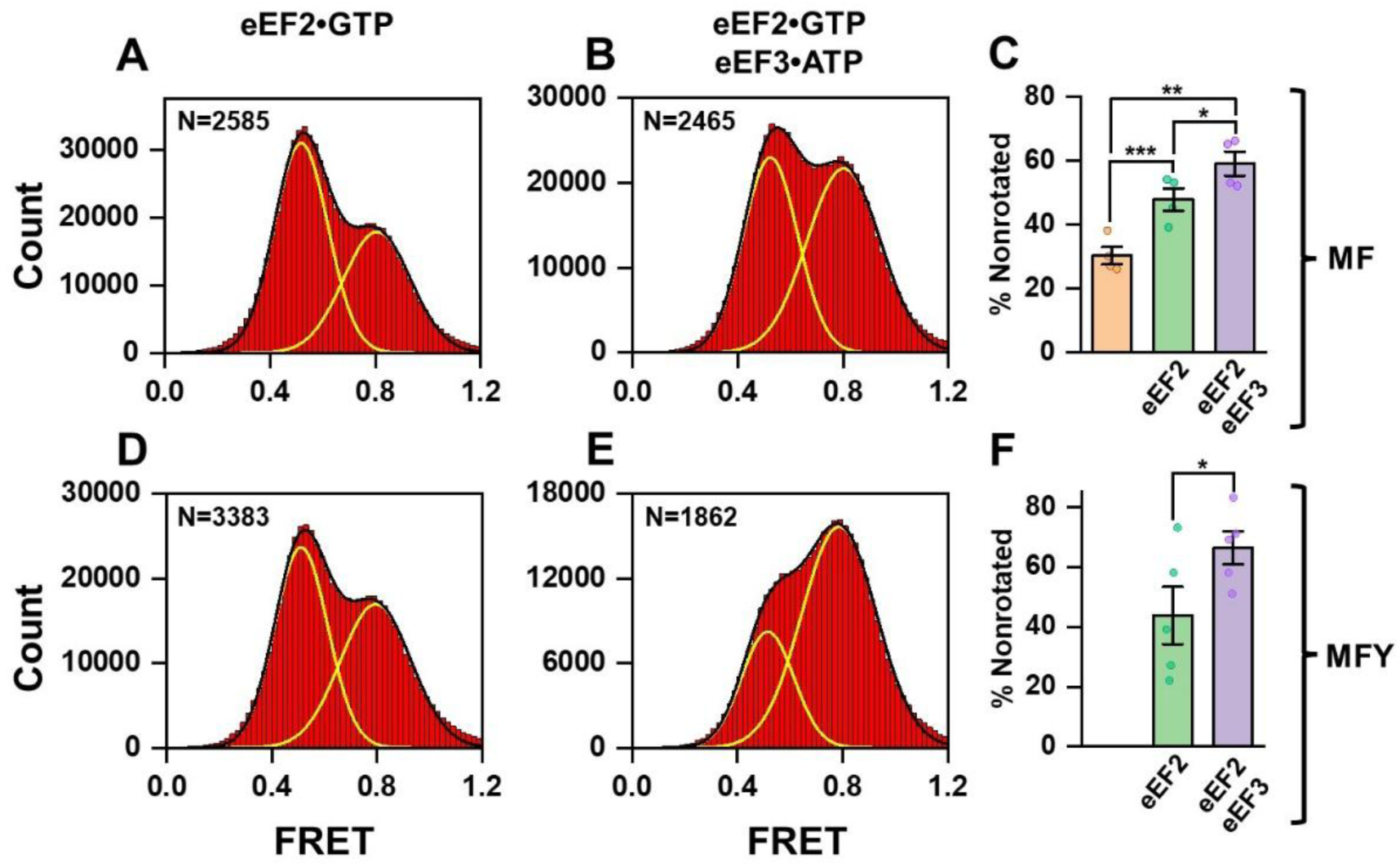
eEF3 enhances eEF2-induced translocation in yeast ribosomes. Pretranslocation ribosomes, which were bound with MFY mRNA, A-site N-Ac-Met-Phe-tRNA^Phe^ and P-site deacylated tRNA^Met^, were incubated for 10 minutes with 1 μM eEF2 and 0.5 mM GTP alone (A, D) or together with 1 μM eEF3 and 0.5 mM ATP (B, E) and then imaged (please see schematics in Supplementary Fig. S5). To enable two elongation cycles (synthesis of MFY tri-peptide followed by translocation), Tyr-tRNA^Tyr^•eEF1A•GTP was included in addition to eEF2/eEF3 (D-F) (please see schematics in Supplementary Fig. S5). Histograms (A-B, D-E) show FRET distribution in uS15-Cy3/eL30-Cy5 ribosomes. Yellow and black lines indicate individual Gaussian fits and the sum of Gaussian fits, respectively. N indicates the total number of FRET traces compiled into each histogram from 4-5 experimental replicates. Bar graphs (C, F) show the fraction of ribosomes in the 0.8 FRET (NR) state observed in pretranslocation ribosomes in the absence of eEF2/eEF3 (salmon), incubated with eEF2•GTP (green) or eEF2•GTP and eEF3•ATP (purple). P-values indicated on the graphs were calculated with a one-tailed heteroscedastic Student t-test, and error bars represent the standard error of the mean (SEM) calculated from 4-5 replicates.

To enable another elongation cycle and synthesis of the tripeptide Met-Phe-Tyr, we incubated uS15-Cy3/eL30-Cy5 80S ribosomes, which contained A-site N-Ac-Met-Phe-tRNA^Phe^ and P-site deacylated tRNA^Met^, with eEF2•GTP and eEF1A•GTP•Tyr-tRNA^Tyr^. The efficiency of eEF2-induced translocation during the second elongation cycle appeared low, as only ∼40% of resulting complexes were observed in the nonrotated (0.8 FRET) posttranslocation conformation (Fig. 6D, F). By contrast, after incubating uS15-Cy3/eL30-Cy5 80S ribosomes, which contained A-site N-Ac-Met-Phe-tRNA^Phe^ and P-site deacylated tRNA^Met^, with eEF2•GTP, eEF1A•GTP•Tyr-tRNA^Tyr^ and eEF3•ATP, ∼70% of ribosomes exhibited 0.8 FRET, indicating more effective translocation linked to switching into the nonrotated conformation (Fig. 6E, F). Similar eEF3-mediated increases in the fraction of the nonrotated conformation in ribosomes incubated with eEF2 was observed when pretranslocation complexes were assembled with leaderless MFV mRNA (Supplementary Fig. S6). A shift in the effect of eEF3 on the extent of translocation, which is manifested by the increase of ribosomes in the nonrotated conformation during the synthesis of a tripeptide, is consistent with data indicating that eEF3 is essential for the second and following elongation cycles (28).

We next tested how eEF3 affects the rate of reverse intersubunit rotation during eEF2-induced translocation. To that end, at 10 s of imaging, eEF2•GTP with or without eEF3•ATP was injected into a microscope slide bound with pretranslocation uS15-Cy3/eL30-Cy5 80S ribosomes, which contained A-site N-Ac-Met-Phe-tRNA^Phe^ and P-site deacylated tRNA^Met^. The dwell time τ_trl(translocation)_ between the injection and the transition from 0.5 FRET to 0.8 FRET reflects the translocation rate (Fig. 7A, C). Switching from 0.5 to 0.8 FRET in the presence of eEF2•GTP alone (Fig. 7A-B) and eEF2•GTP/eEF3•ATP (Fig. 7C-D) occurred at rates 0.5 and 0.4 s^-1^, respectively. These rates are faster than the translocation rate (0.15 s^-1^ – 0.1 min^-1^) measured in previous *in vitro* experiments performed with yeast ribosomes (28), but substantially slower than the 5-10 amino acids per second rates reported for protein synthesis in live yeast cells (61,62). This is likely due to higher *in vivo* concentrations of elongation factors than the 1 μM concentrations of eEF2 and eEF3 used in our experiments. Other biochemical factors, such as His-tagging of eEF2 and eEF3, and performing all kinetic smFRET experiments at room temperature instead of the 30°C optimal for yeast growth, may contribute to slowing translocation rates in the *in vitro* experiments. Nevertheless, our experiments clearly indicate that eEF3 does not accelerate the rate of eEF2-induced reverse intersubunit rotation. Our results are consistent with the model, suggesting that eEF3 facilitates E-site tRNA release at the late step of translocation (28), which follows eEF2-induced reverse intersubunit rotation and is required to enable the next elongation cycle.

**Figure 7.**
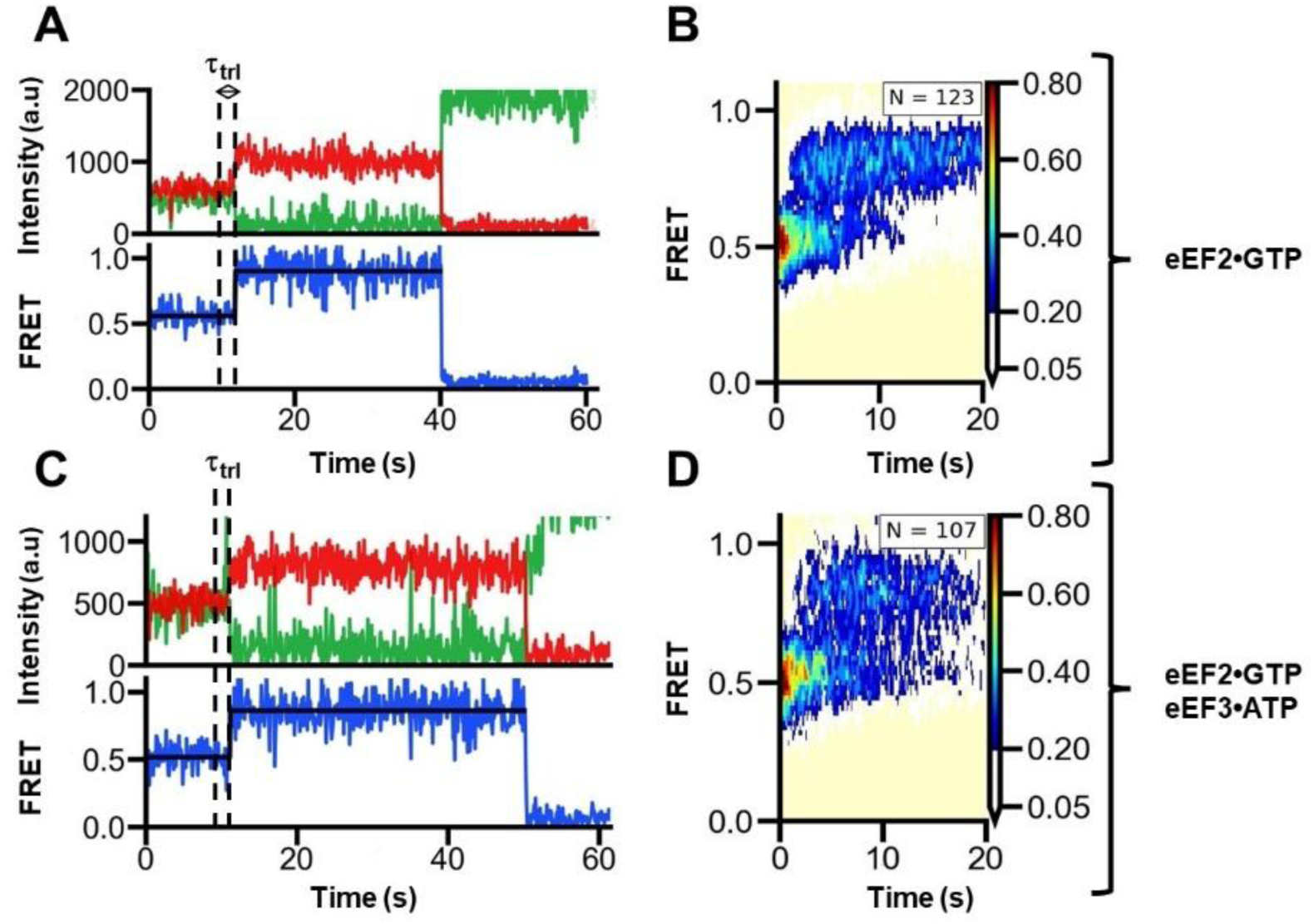
eEF3 does not affect the rate of eEF2-induced reverse intersubunit rotation. During imaging of pretranslocation uS15-Cy3/eL30-Cy5 ribosomes, which were programmed with MFY mRNA and bound with A-site N-Ac-Met-Phe-tRNA^Phe^ and P-site deacylated tRNA^Met^, mRNA-tRNA translocation was induced by injecting eEF2•GTP alone (A-B) or eEF2•GTP/eEF3•ATP (C-D) into the microscope slide at the 10 s mark. (A, C) Representative FRET traces show donor fluorescence (green), acceptor fluorescence (red), and FRET efficiency (blue). The two-state hidden Markov model fit is shown by the black line. τ_trl_ represents the dwell time between the injection of eEF2 and the R-to-NR transition. (B, D) Surface contour plots were generated by superimposition of smFRET traces post-synchronized at the time of eEF2 or eEF2-eEF3 injection. N is the number of traces combined.

To test whether eEF3•ATP alone (i.e., without eEF2) can promote translocation, we incubated pretranslocation uS15-Cy3/eL30-Cy5 80S ribosomes, which contained A-site N-Ac-Met-Phe-tRNA^Phe^ and P-site deacylated tRNA^Met^, with eEF3•ATP for 10 minutes at 30°C before removing eEF3•ATP from the slide and imaging. Incubation with eEF3•ATP without eEF2 did not noticeably alter the ratio of NR and R conformation in the ribosome population (Fig. 8A). In contrast, an increase in the NR population was observed when pretranslocation ribosomes were simultaneously treated with both eEF2•GTP and eEF3•ATP (Fig, 6B-C), or sequentially, first with eEF3•ATP alone and then with both eEF2•GTP and eEF3•ATP before removing translation factors from the slide and imaging (Supplementary Fig. S7). Hence, our data do not provide supporting evidence that eEF3 can induce translocation in the absence of eEF2.

**Figure 8.**
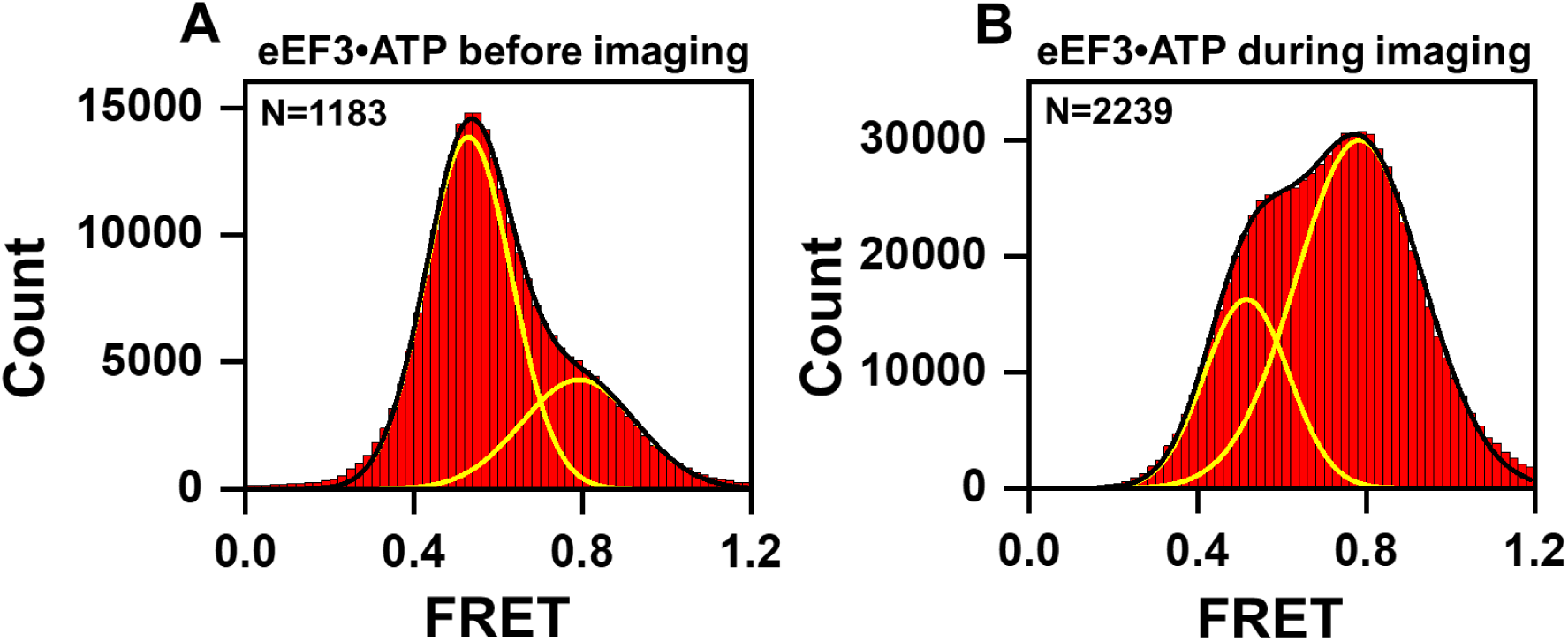
While eEF3•ATP does not induce translocation in the absence of eEF2, eEF3•ATP binding stabilizes the NR conformation. Histograms (A-B) show FRET distribution in uS15-Cy3/eL30-Cy5 ribosomes. (A) Pretranslocation ribosomes, which were bound with MFY mRNA, A-site N-Ac-Met-Phe-tRNA^Phe^ and P-site deacylated tRNA^Met^, were pre-incubated at 30°C for 10 minutes with 1 μM eEF3 and 0.5 mM ATP. Then ribosomes were immobilized on the slide and imaged in the absence of eEF3•ATP. (B) Pretranslocation ribosomes were imaged in the presence of 1 μM eEF3 and 0.5 mM ATP. Yellow and black lines indicate individual Gaussian fits and the sum of Gaussian fits, respectively. N indicates the number of FRET traces compiled into the histogram.

After establishing that incubating pretranslocation ribosomes with eEF3•ATP and then removing eEF3•ATP does not result in appreciable changes in the ratio of R vs NR conformations, we next tested whether stable binding of eEF3 favors the nonrotated conformation of the ribosome, as was suggested by previous cryo-EM studies (26,28). How ATP hydrolysis and eEF3 release from the ribosome are regulated is unclear. Nevertheless, published biochemical data show that eEF3 tightly binds to the ribosome in the ATP-bound conformation, either with ATP or non-hydrolysable ATP analogues, while binding in the presence of ADP is less stable (26). Consistent with ATP being required for binding, the R vs NR ratio of pretranslocation ribosomes remained unaltered when they were imaged in the presence of 1 μM eEF3 without an added nucleotide (Supplementary Fig. S8A). Surprisingly, 1 μM eEF3 combined with 0.5 mM nonhydrolyzable ATP analogue, ADPNP, did not affect the R vs NR ratio of pretranslocation ribosomes when incubated during imaging either (Supplementary Fig. S8B). In contrast, when pretranslocation ribosomes were imaged with 1 μM eEF3 and 0.5 mM ATP, the fraction of 0.8 FRET increased from 35 to 70% (Fig. 8B), indicating stabilization of NR conformation of the ribosome. It is possible that subtle structural differences between ADPNP and ATP, such as differences in charge geometry and distribution, explain the inability of eEF3•ADPNP to appreciably alter the R vs NR equilibrium. Consistent with this hypothesis, obtaining cryo-EM structures of the eEF3•ADPNP•ribosome complex required adding covalent cross-linking agents such as glutaraldehyde (28).

Cryo-EM reconstructions of eEF3-ribosome complexes revealed that eEF3 bridges the ribosomal subunits in the nonrotated conformation, as the N- and C-terminal domains of eEF3 interact with the 40S and 60S subunits, respectively (26,28). Hence, eEF3 likely stabilizes the NR conformation of the ribosome by slowing the NR-to-R transition. However, we could not test this hypothesis in our smFRET data because only ∼2% of traces showed transitions between 0.5 and 0.8 FRET states in pretranslocation ribosomes imaged with or without eEF3. The observations that eEF3 and eEF2 favor two different conformations of the ribosome (Fig. 8, Fig. 6) (17,18,28), namely NR and R, are consistent with the model suggesting that eEF3 is not involved in the eEF2-mediated steps of translocation, such as reverse intersubunit rotation (Fig. 7). Taken together, our equilibrium and kinetic smFRET experiments support the model that eEF3-induced ejection of deacylated tRNA from the E site occurs at the late step of translocation and is required to enable the ribosome to undergo the next elongation cycle.

### uS15-Cy3/eL30-Cy5 FRET as a tool for following eukaryotic translation elongation

Our uS15-Cy3/eL30-Cy5 FRET data show that yeast ribosomes switch between two predominant conformations corresponding to 0.8 and 0.5 FRET states (Fig. 4). Observed FRET changes are consistent with ∼10° intersubunit rotation between the nonrotated (NR) and fully rotated (R) conformations of the ribosome, in which tRNAs are bound in the classical and hybrid states, respectively. Indeed, hindering binding of deacylated P-site tRNA in the P/E hybrid state by cycloheximide reduced the fraction of ribosomes observed in 0.5 FRET state (Fig. 5). Similar to bacterial ribosomes, the A-site tRNA binding/transpeptidation and mRNA-tRNA translocation steps of the elongation cycle in eukaryotic ribosomes appear to be coupled to NR-to-R and R-to-NR transitions (Fig. 2, 3, 6-7, Supplementary Fig. S4, 6).

Our uS15-Cy3/eL30-Cy5 FRET data did not demonstrate sampling of additional FRET states besides 0.8 and 0.5 FRET. Only transitions between 0.5 and 0.8 FRET were observed in traces with FRET fluctuations (Fig. 4C, 7A, C). Evidently, uS15-Cy3/eL30-Cy5 FRET is not sensitive to either subtle (∼4°) partial intersubunit rotation or the subunit rolling seen in steady-state cryo-EM reconstructions of yeast ribosomes (5,49) because these rearrangements lead to modest changes in the distance between fluorophores. Indeed, between cryo-EM structures of the partially rotated, “TI-2” conformation (PDB 8CDR) and nonrotated, posttranslocation (PDB 8CGN) state of the yeast ribosome (49), distances between C termini of uS15 and eL30 differed by only 1.5 Å.

Our data indicate that the uS15-Cy3/eL30-Cy5 FRET pair serves as a robust and reliable tool for following rearrangements between the NR and R conformations of eukaryotic ribosome during translation elongation. Hence, this single-molecule assay can be employed for future investigation of co-translational regulatory events in eukaryotic ribosomes such as ribosome pausing, stalling, collision, and frameshifting, or the study of antibiotic binding and action.

## Resource availability

Further information and requests for resources should be directed to and will be fulfilled by the lead contact, Dmitri Ermolenko.

## Materials availability

Yeast strains generated in this study are available upon request.

## ACKNOWLEDGEMENTS

We thank Eric Phizicky and Elizabeth Grayhack for providing the BCY123 strain of *S. cerevisiae* and advice on yeast genetics, Anton Komar for advice on yeast ribosome purification, Alexei Petrov for providing the eEF1Bα expression plasmid, Terri Kinzy and Andrei Korostelev for the 6His-eEF2 and 6His-eEF3 expression yeast strains, Ruben Gonzalez for sharing the tMAVEN software before its publication, Riley Gentry for advice on using the tMAVEN and vbscope softwares. This work was supported by National Institutes of Health grant R35GM141812 (to D.N.E). A.K.G. was supported by National Institutes of Health grant T32-GM145461. Phosphoimaging was done using Typhoon RGB instrument, the purchase of which was supported by NIH Equipment Grant (S10-OD021489-01A1).

## AUTHORS CONTRIBUTIONS

D.N.E. conceived the project. H.W. performed genetic manipulations as well as preliminary smFRET and biochemical experiments. A.V. I. and A.K.G. performed biochemical preparative experiments, A.D. and A.K.G. performed smFRET experiments and analyzed smFRET data. D.N.E. and A.K.G. wrote the original draft of the manuscript. All authors contributed to manuscript writing and editing.

