## Supplementary Materials for "Observing intersubunit dynamics in single yeast ribosomes"

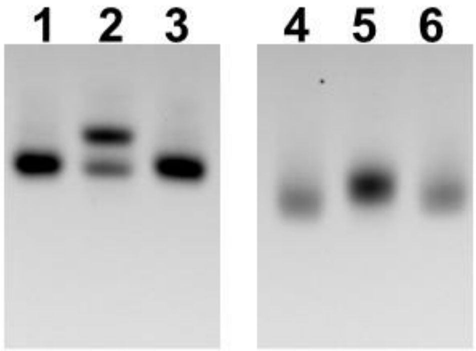


**Supplementary Figure S1.** The extent of aminoacylation of yeast *S. cerevisiae* $tRNA^{Phe}$ and bacterial E.coli $tRNA^{Met}$ was revealed by denaturing acid PAGE. Deacylated $tRNA^{Phe}$ – lanes 1 and 3, Phe-$tRNA^{Phe}$ – lane 2, deacylated $tRNA^{Met}$ – lanes 4 and 6, N-Ac-Met-$tRNA^{Met}$ – lane 5. Upward-shifted bands in lanes 2 and 5 correspond to aminoacylated tRNA species.

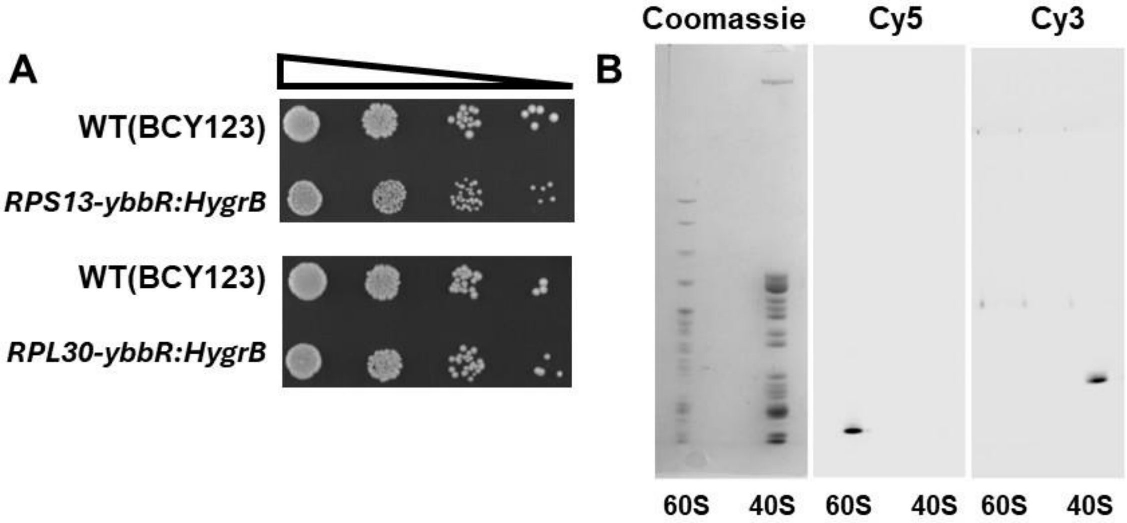


**Supplementary Figure S2.** Introducing Cy3 and Cy5 fluorophores into proteins uS15 and eL30. (A) Effects of introducing ybbR tag into proteins uS15 (RPS13) and eL30 (RPL30) on cell growth. Strains were grown on YPD plates for 48 hours at 30°C. (B) Proteins of 40S-uS15-Cy3 and 60S-eL30-Cy5 subunits were analyzed using SDS-PAGE. Proteins were visualized in the gel by staining the gel with Coomassie brilliant blue dye, exciting Cy5 fluorescence, or exciting Cy3 fluorescence. Single Cy3 and Cy5 fluorescent bands indicate specific labeling of uS15 and eL30 proteins, respectively.

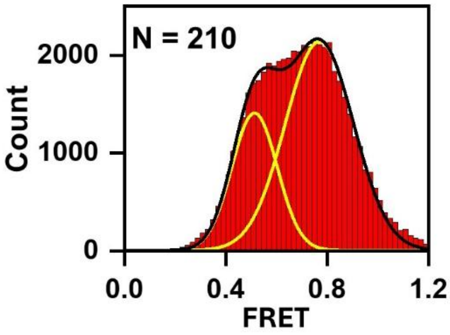


**Supplementary Figure S3.** GTP is necessary for eEF1A-mediated tRNA binding. Ribosomes programmed with MFY mRNA and bound with P-site N-Ac-Met-tRNA$^{Met}$ were incubated with eEF1A and Phe-tRNA$^{Phe}$ in the absence of GTP and then imaged. Yellow and black lines indicate individual Gaussian fits and the sum of Gaussian fits, respectively. N indicates the number of FRET traces compiled into the histogram.

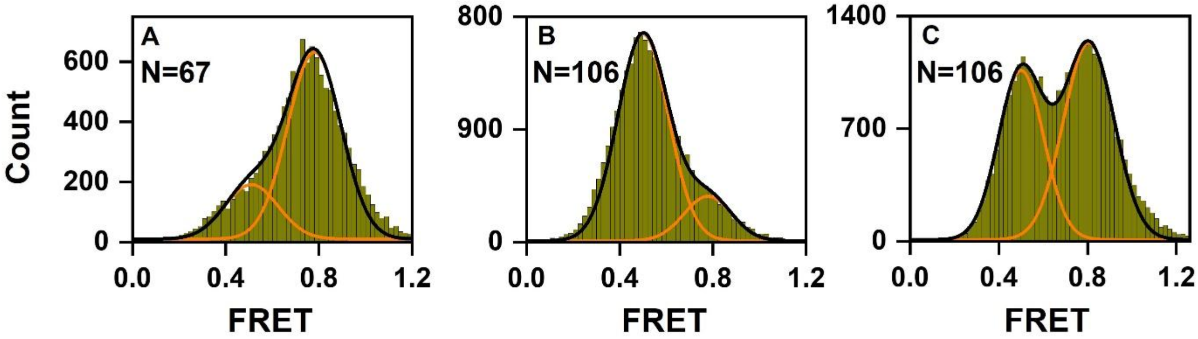


**Supplementary Figure S4.** Assembly of ribosomal complexes from 40S-uS15-Cy3 and 60S-eL30-Cy5 subunits. Histograms (A-C) show FRET distribution in uS15-Cy3/eL30-Cy5 ribosomes. 40S-uS15-Cy3 and 60S-eL30-Cy5 subunits were incubated with N-Ac-Met-tRNA$^{Met}$ and MFY mRNA. Resulting ribosomal complexes containing P-site N-Ac-Met-tRNA$^{Met}$ (A) were treated with eEF1A•GTP•Phe-tRNA$^{Phe}$ (B) and then eEF2•GTP (C). Yellow lines indicate individual Gaussian fits. Black line shows the sum of Gaussian fits. N indicates the number of FRET traces compiled into each histogram.

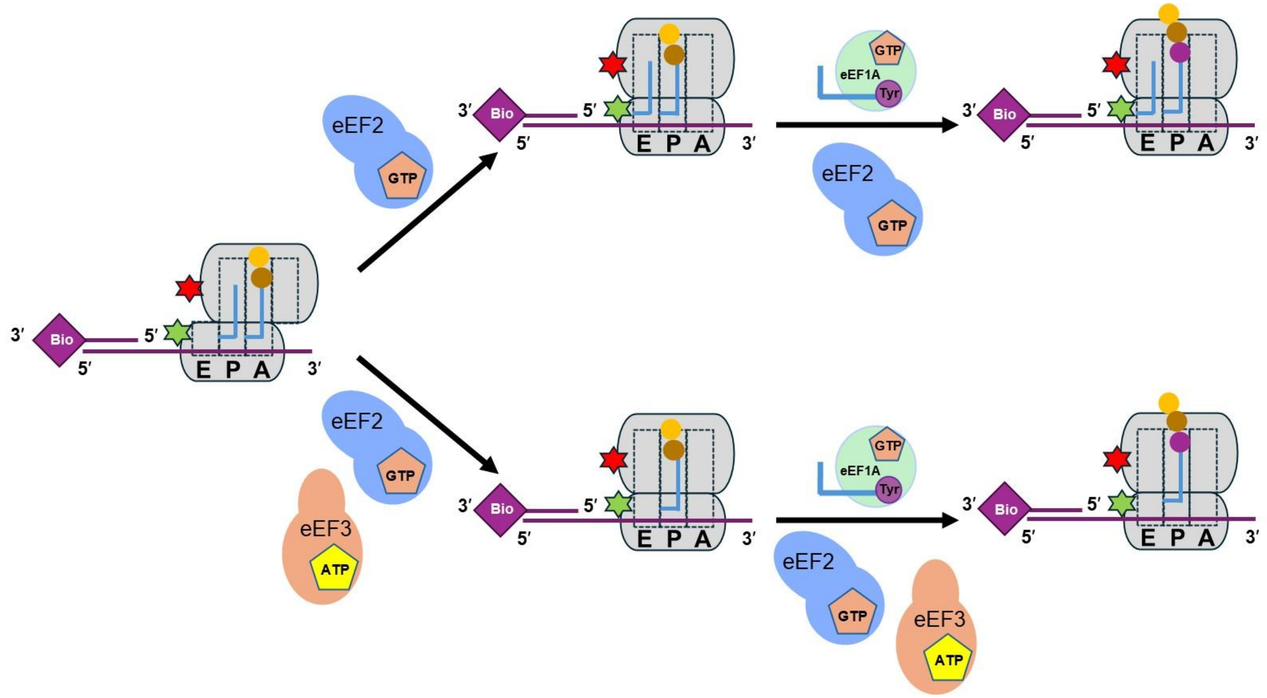


**Supplementary Figure S5.** eEF2 or eEF2/eEF3 induce tRNA translocation. Pre-translocation ribosomes (left) programmed with MFY mRNA and bound to A-site N-Ac-Met-Phe-tRNA$^{Phe}$ and P-site deacylated tRNA$^{Met}$ were pre-incubated at 30°C for 10 minutes with 1 μM eEF2 and 0.5 mM GTP without (Fig. 6 A, C) or with 1 μM eEF3 and 0.5 mM ATP (Fig. 6 B-C). To enable two elongation cycles (synthesis of MFY tri-peptide followed by translocation), Tyr-tRNA$^{Tyr}$•eEF1A•GTP was included in addition to eEF2/eEF3 (Fig. 6 D-F).

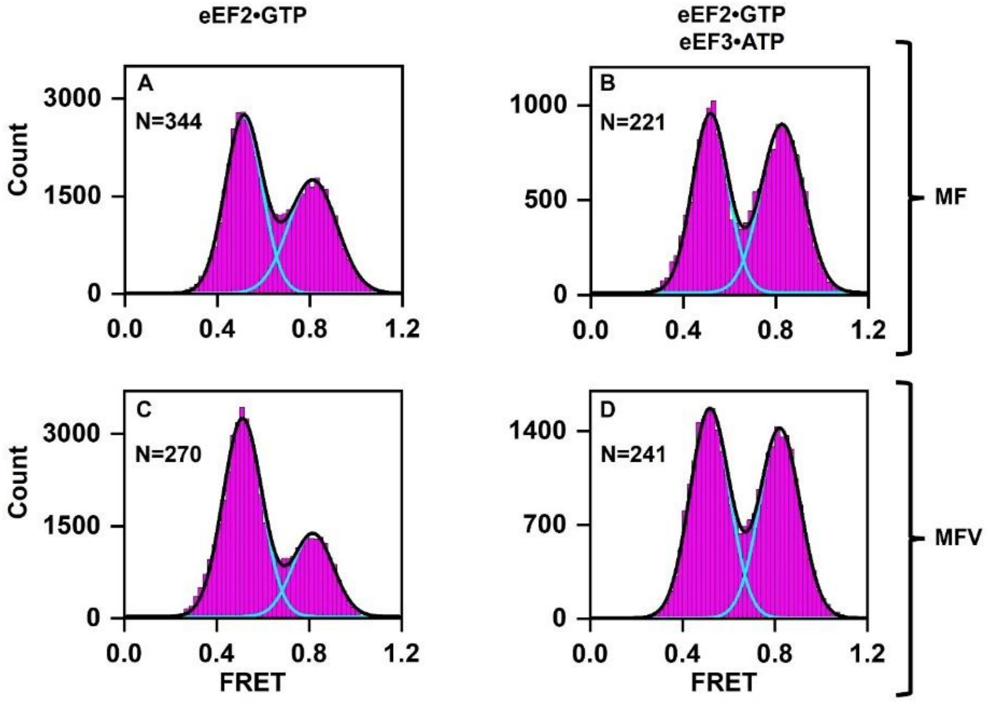


**Supplementary Figure S6.** eEF3 enhances eEF2-induced translocation in yeast ribosomes programmed with MFV mRNA tethered to the slide at the 3' end. Histograms (A-D) show FRET distribution in uS15-Cy3/eL30-Cy5 ribosomes. Pre-translocation ribosomes, which were bound with MFV mRNA, A-site N-Ac-Met-Phe-tRNA$^{Phe}$ and P-site deacylated tRNA$^{Met}$, were incubated for 10 minutes at 30°C with eEF2•GTP alone (A, C) or eEF2•GTP/eEF3•ATP (B, D) and then imaged. To enable two elongation cycles (i.e. synthesis of MFV tri-peptide followed by translocation), Val-tRNA$^{Val}$•eEF1A•GTP was included in addition to eEF2/eEF3 (C-D). mRNA-tRNA translocation results in switching from the R (0.5 FRET) to NR (0.8 FRET) conformation. Cyan and black lines indicate individual Gaussian fits and the sum of Gaussian fits, respectively. N indicates the number of FRET traces compiled into each histogram.

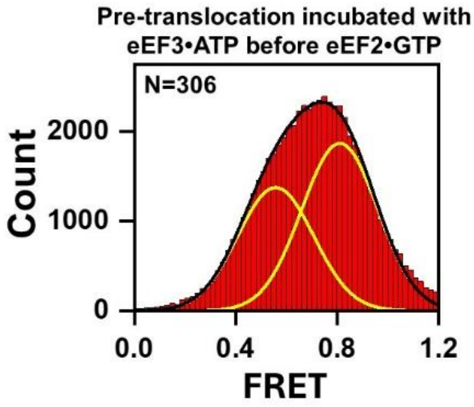


**Supplementary Figure S7.** The order of eEF2/eEF3 pre-incubation does not influence population dynamics. Pre-translocation ribosomes, which were bound with MFY mRNA, A-site N-Ac-Met-Phe-tRNA$^{Phe}$ and P-site deacylated tRNA$^{Met}$, were pre-incubated at 30°C for 10 minutes with eEF3•ATP, then pre-incubated an additional 10 minutes at 30°C with eEF2•GTP, and then imaged. Yellow and black lines indicate individual Gaussian fits and the sum of Gaussian fits, respectively. N indicates the number of FRET traces compiled into the histogram.

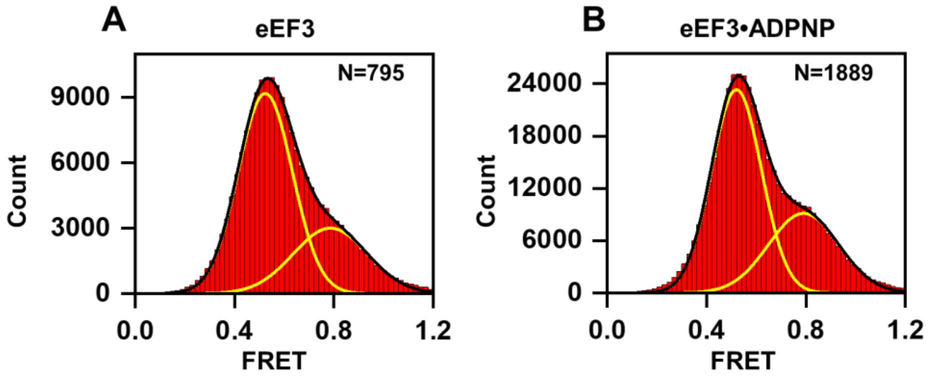


**Supplementary Figure S8.** eEF3 does not influence the conformation of the pre-translocation ribosome without ATP. Histograms (A-B) show FRET distribution in uS15-Cy3/eL30-Cy5 ribosomes. Pre-translocation ribosomes, which were bound with MFY mRNA, A-site N-Ac-Met-Phe-$tRNA^{Phe}$ and P-site deacylated $tRNA^{Met}$, were incubated for 1 μM eEF3 alone (A) or eEF3•ADPNP (B) in the imaging buffer during data collection. Yellow and black lines indicate individual Gaussian fits and the sum of Gaussian fits, respectively. N indicates the number of FRET traces compiled into each histogram.

| Fig. Panel | mRNA | tRNAs | Elongation Factors, Nucleotides & Antibiotics | Low FRET peak center | Low FRET % area | High FRET peak center | High FRET % area |
|---|---|---|---|---|---|---|---|
| 2A | MFV | NAcMet-tRNA$^{Met}$ | | 0.55 | 21 | 0.80 | 79 |
| 2B | MFV | NAcMet-tRNA$^{Met}$, Phe-tRNA$^{Phe}$ | eEF1A•GTP | 0.51 | 76 | 0.82 | 24 |
| 2C | MFY | NAcMet-tRNA$^{Met}$ | | 0.52 | 18 | 0.79 | 82 |
| 2D | MFY | NAcMet-tRNA$^{Met}$, Phe-tRNA$^{Phe}$ | eEF1A•GTP | 0.51 | 65 | 0.78 | 35 |
| 3A | MFY | NAcMet-tRNA$^{Met}$, Phe-tRNA$^{Phe}$ | eEF1A•GTP<br>20 μM didemnin B | 0.51 | 27 | 0.79 | 73 |
| 3B | MFY | NAcMet-tRNA$^{Met}$, Phe-tRNA$^{Phe}$ | eEF1A•GTP<br>70 μM didemnin B | 0.51 | 13 | 0.82 | 87 |
| 5A | MFY | NAcMet-tRNA$^{Met}$, Phe-tRNA$^{Phe}$ | eEF1A•GTP | 0.53 | 71 | 0.80 | 29 |
| 5B | MFY | NAcMet-tRNA$^{Met}$, Phe-tRNA$^{Phe}$ | eEF1A•GTP<br>5 μM CHX | 0.54 | 60 | 0.79 | 40 |
| 5C | MFY | NAcMet-tRNA$^{Met}$, Phe-tRNA$^{Phe}$ | eEF1A•GTP<br>50 μM CHX | 0.53 | 31 | 0.77 | 69 |
| 5D | MFY | NAcMet-tRNA$^{Met}$, Phe-tRNA$^{Phe}$ | eEF1A•GTP<br>500 μM CHX | 0.53 | 30 | 0.77 | 70 |
| 6A | MFY | NAcMet-tRNA$^{Met}$, Phe-tRNA$^{Phe}$ | eEF1A•GTP,<br>eEF2•GTP | 0.51 | 50 | 0.79 | 50 |
| 6B | MFY | NAcMet-tRNA$^{Met}$, Phe-tRNA$^{Phe}$ | eEF1A•GTP,<br>eEF2•GTP,<br>eEF3•ATP | 0.52 | 44 | 0.80 | 56 |
| 6D | MFY | NAcMet-tRNA$^{Met}$, Phe-tRNA$^{Phe}$, Tyr-tRNA$^{Tyr}$ | eEF1A•GTP,<br>eEF2•GTP | 0.51 | 51 | 0.80 | 49 |
| 6E | MFY | NAcMet-tRNA$^{Met}$, Phe-tRNA$^{Phe}$, Tyr-tRNA$^{Tyr}$ | eEF1A•GTP,<br>eEF2•GTP,<br>eEF3•ATP | 0.52 | 27 | 0.79 | 73 |
| 8A | MFY | NAcMet-tRNA$^{Met}$, Phe-tRNA$^{Phe}$ | eEF1A•GTP,<br>eEF3•ATP | 0.53 | 71 | 0.79 | 29 |
| 8B | MFY | NAcMet-tRNA$^{Met}$, Phe-tRNA$^{Phe}$ | eEF1A•GTP,<br>eEF3•ATP* | 0.52 | 27 | 0.78 | 73 |
| S3 | MFY | NAcMet-tRNA$^{Met}$, Phe-tRNA$^{Phe}$ | eEF1A | 0.51 | 30 | 0.77 | 70 |
| S4A | MFY | NAcMet-tRNA$^{Met}$ | | 0.51 | 23 | 0.78 | 77 |
| S4B | MFY | NAcMet-tRNA$^{Met}$, Phe-tRNA$^{Phe}$ | eEF1A•GTP | 0.50 | 83 | 0.78 | 17 |
| S4C | MFY | NAcMet-tRNA$^{Met}$, Phe-tRNA$^{Phe}$ | eEF1A•GTP,<br>eEF2•GTP | 0.50 | 43 | 0.81 | 57 |
| S6A | MFV | NAcMet-tRNA$^{Met}$, Phe-tRNA$^{Phe}$ | eEF1A•GTP,<br>eEF2•GTP | 0.51 | 55 | 0.81 | 45 |
| S6B | MFV | NAcMet-tRNA$^{Met}$, Phe-tRNA$^{Phe}$ | eEF1A•GTP,<br>eEF2•GTP,<br>eEF3•ATP | 0.52 | 48 | 0.83 | 52 |
| S6C | MFV | NAcMet-tRNA$^{Met}$, Phe-tRNA$^{Phe}$, Val-tRNA$^{Val}$ | eEF1A•GTP,<br>eEF2•GTP | 0.51 | 70 | 0.82 | 30 |
| S6D | MFV | NAcMet-tRNA$^{Met}$, Phe-tRNA$^{Phe}$, Val-tRNA$^{Val}$ | eEF1A•GTP,<br>eEF2•GTP,<br>eEF3•ATP | 0.52 | 52 | 0.82 | 48 |

| | | | | | | | |
|---|---|---|---|---|---|---|---|
| S7 | MFY | NAcMet-tRNA$^{Met}$, Phe-tRNA$^{Phe}$ | eEF1A•GTP, eEF2•GTP eEF3•ATP | 0.56 | 42 | 0.81 | 58 |
| S8A | MFY | NAcMet-tRNA$^{Met}$, Phe-tRNA$^{Phe}$ | eEF1A•GTP, eEF3* | 0.52 | 69 | 0.79 | 31 |
| S8B | MFY | NAcMet-tRNA$^{Met}$, Phe-tRNA$^{Phe}$ | eEF1A•GTP, eEF3•ADPNP* | 0.52 | 65 | 0.79 | 35 |

**Supplementary Table S1.** Composition of assembled complexes and Gaussian fitting parameters. mRNAs, tRNAs, antibiotics, and elongation factors added during each complex assembly are indicated. The asterisk * designates experiments in which elongation factors were present in the slide during imaging (i.e. in the imaging buffer) rather than pre-incubated and removed from the slide prior to data collection. smFRET traces collected from each experiment were extracted and FRET efficiencies were compiled into histograms that were fit to two Gaussian distributions centered around the low FRET state (average 0.52, S.D. 0.01) and high FRET state (average 0.80, S.D. 0.02). The area under each curve was used to calculate the percent occupancy of the low and high FRET states for each complex.
